# Endothelin-converting enzyme 2 differentially regulates kappa opioid receptor trafficking and function

**DOI:** 10.1101/2025.03.11.642672

**Authors:** Achla Gupta, Ivone Gomes, Salvador Sierra, Aya Osman, Lakshmi A. Devi

## Abstract

Following activation by endogenous opioid peptides, mu and delta opioid receptors have been shown to undergo differential internalization and recycling; the rate and extent of recycling but not internalization was found to be regulated by the endocytic peptide converting enzyme, ECE2. This study focuses on kappa opioid receptors (KOR) and the ability of endogenous opioid peptides released by post-translational processing of prodynorphin and proenkephalin to induce KOR and ECE2 internalization/recycling, and how ECE2 modulates these processes. Using a proximity-based ligation assay we show that KOR and ECE2 are in close proximity to facilitate co-internalization. In addition, we find that treatment with longer opioid peptides induces fast and robust internalization and recycling of ECE2 at a rate and extent comparable to that of KOR. Next, we directly examined the role of ECE2 in modulating KOR recycling. We find that ECE2 inhibition significantly attenuates KOR recycling. Finally, we examined the role of ECE2 in modulating KOR signaling and find that resensitization of KOR by peptides that are substrates of ECE2 are attenuated by ECE2 inhibition. Taken with the differential expression of ECE2 in the brain (relatively high expression in midbrain & hypothalamus and low expression in the striatum & hippocampus), these results highlight a pivotal role for ECE2 in differentially modulating KOR function.

**Significance Statement:** This study highlights a role for ECE2 in agonist mediated regulation of KOR function by select prodynorphin and proenkephalin derived peptides. Collectively our studies suggest that ECE2 inhibitors could be developed as therapeutics for pathologies involving dysregulations in KOR signaling.

## Introduction

The endogenous opioid system plays a role in regulating a wide range of biological processes and consists of three opioid receptors, mu (MOR), delta (DOR), and kappa (KOR) and peptides acting at these receptors (Stein, 2016). Endogenous opioid peptides are generated from the proteolytic cleavage of the precursor proteins prodynorphin (proDYN), proenkephalin (proENK) and proopiomelanocortin (POMC), giving rise to >20 peptides (Stein, 2016; Shenoy and Lui, 2023). Of these, proENK-derived peptides are the most abundant (Fricker, 2012) and previously thought to exclusively activate DOR and MOR while proDYN-derived peptides are less abundant and thought to selectively bind to and activate KOR (Pasternak, 2011). However, mounting evidence suggests that all endogenous opioid peptides display differential activity at the three opioid receptors, with the biological significance of this phenomena still being understood (Gomes et al., 2020).

In addition to G protein-dependent signaling, peptide activation of opioid receptors can initiate G protein-independent signaling (Gomes et al., 2020) following β-arrestin recruitment to the phosphorylated receptor, where it functions as a scaffold enabling signaling through different molecules (Luttrell and Lefkowitz, 2002; DeFea, 2008). These two pathways of signaling can be differentially activated by receptor agonists including opioid peptides, known as biased agonism (Gomes et al., 2020). Following β-arrestin recruitment, the receptor is internalized and targeted for either recycling or degradation (von Zastrow and Williams, 2012), a process known as receptor trafficking. While the mechanisms underlying opioid receptor internalization and desensitization have been well studied (Hanyaloglu and Von Zastrow, 2008), fewer studies have assessed receptor recycling and resensitization mechanisms. Understanding and having the ability to modify the molecular machinery involved in recycling and resensitization is important as these processes ultimately modulate receptor signaling and are of importance in diseases involving dysregulations in receptor signaling such as addiction, chronic pain and cancer.

During acute treatment with a ligand, the receptor endocytosed with its peptide ligand is thought to dissociate from β-arrestin and the bound peptide in an acidic endocytic compartment leading to receptor dephosphorylation and recycling to the cell surface and/or trafficking to a degradative compartment (Williams et al., 2013). The endocytic fate of activated receptors has been shown to be ligand dependent (Kunselman et al., 2021), where the peptide ligand endocytosed along with the receptor could be selectively processed to further modulate the rate and extent of receptor recycling and subsequent signaling (Mellman et al., 1986) although not much is known about the proteolytic enzymes involved.

In this context, Endothelin Converting Enzyme 2 (ECE2), a member of the metalloprotease family exhibits endocytic localization, optimum activity at acidic pH and plays a role in non-classical hydrolysis of opioid peptide substrates in an acidic endocytic compartment (Emoto and Yanagisawa, 1995; Mzhavia et al., 2003; Devi and Miller, 2013). We previously used a selective ECE2 inhibitor, S13649 (Gagnidze et al., 2008) to show that inhibiting ECE2 activity significantly impaired MOR and DOR recycling and subsequent signaling by specific peptide substrates (Gupta et al., 2014, 2015). Although these intriguing findings highlight a role for endosomal peptide processing in shaping receptor function, ECE2 localization in relation to opioid receptors and their peptide ligands remains unknown. Other metalloproteases are expressed either exclusively at the cell surface (eg., neprylisin; NEP) or both at the cell surface and intracellular compartments (eg., Endothelin Converting Enzyme 1; ECE1) (Roosterman et al., 2007). Based on ECE1 we hypothesized that ECE2 is expressed at the cell surface and is internalized with the activated receptor-peptide ligand complex to an acidic endocytic compartment where it selectively degrades peptide substrates, before being recycled back to the cell surface with the receptor.

In this study using KOR as a model system, we addressed the localization of ECE2 using a proximity ligation assay and find that KOR and ECE2 are in close proximity at the cell surface which facilitates co-internalization upon treatment with a peptide ligand. In addition, we find that longer opioid peptide agonists released by post-translational processing of proDYN and proENK induce fast and robust ECE2 internalization and recycling, with internalization and recycling kinetics mirroring those of KOR. Importantly, we find that ECE2 inhibition blocks KOR recycling and signaling. Specifically, receptor resensitization by substrate peptide agonists including Dyn B13, a peptide derived from proDYN, and BAM22 a peptide derived from proENK are blocked by ECE2 inhibition. Together these results indicate that ECE2 is trafficked to and from the cell surface upon KOR activation by peptide ligands, to play a role in differentially modulating KOR signaling and highlight ECE2 as a therapeutic target for modulating opioid receptor function in disease states.

## Results

### Distribution of ECE-2, proENK and proDYN mRNA in mouse brain

In a previous study, we found that ECE2 cleaves select opioid peptides derived from proDYN and proENK processing (Mzhavia et al., 2003). For this to occur in the brain, ECE2 would need to be expressed in the same brain regions as opioid peptide precursors giving rise to opioid peptides listed in **Table 1**. To assess the extent of ECE2 co-expression with proDYN or proENK, the Allen Mouse Brain Atlas (mouse.brain-map.org) was utilized to assess expression profiles. In line with previous findings (Rodriguiz et al., 2008), ECE2 mRNA is found to be highly expressed in hypothalamus, midbrain, cerebellum, frontal cortex, pons and the granule layer of the dentate gyrus of the hippocampus (**Fig 1A**). Moderate levels of ECE2 mRNA were found in limbic structures such as amygdala and striatum (**Fig 1A**). Given the high expression of ECE2 in the dentate gyrus of the hippocampus, a key brain region mediating learning and memory (**Fig 1B**), we next examined the expression of proENK and proDYN mRNA in this brain region and found that their expression overlaps with that of ECE2 (**Fig 1B middle and far right panel**). In contrast, the nucleus accumbens (NAc), a key brain region involved in mediating reward, was found to have moderate ECE2 expression levels and high expression of proENK and proDYN mRNA (**Fig 1C**). Together, the overlapping mRNA distribution studies suggest that ECE2 could play a role in processing of peptides derived from proDYN and proENK in specific brain regions, including those associated with learning and memory.

**Fig 1.**
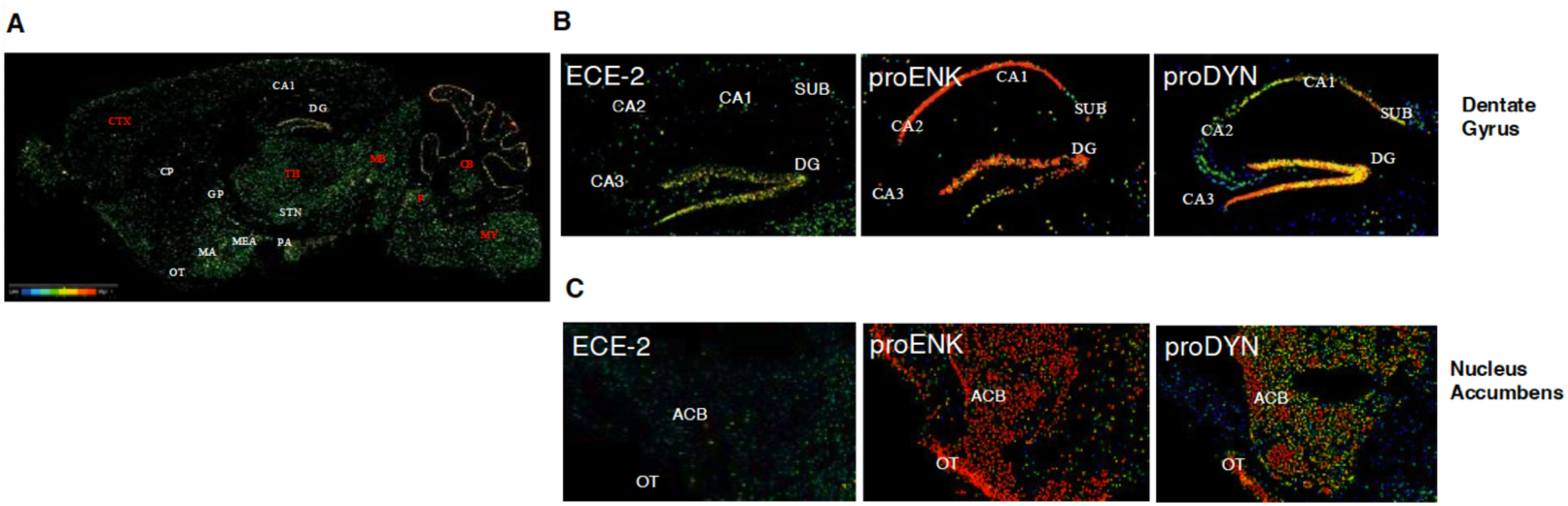
ECE2 mRNA expression overlaps with proENK and proDYN mRNA in brain regions associated with learning and memory. mRNA expression data was obtained from Allen Brain Mouse Atlas (mouse.brain-map.org) and shows in situ hybridization of sagittal mouse brain sections probed with ECE2, proENK and proDYN (**A)** Distribution of ECE2 mRNA in mouse brain **(B)** Distribution of ECE2, proENK and proDYN mRNA in Dentate Gyrus **(C)** Distribution of ECE2, proENK and proDYN mRNA in Nucleus Accumbens. Abbreviations: CTX: Cerebral Cortex CP: Caudate Putamen OT: Olfactory Tubercle TH: Thalamus MY: Medulla CB: Cortex MB: Midbrain DG: Dentate Gyrus P: Pons STN: Subthalamic nucleus GP: Globus Pallidus MA: medial amygdaloid nucleus MEA: medial nucleus of amygdala.

**Table 1.** Endogenous opioid peptides used and their amino acid sequence (human/rat/mouse).

| Precursor | Peptide | Peptide sequence |
| --- | --- | --- |
| Prodynorphin | $\beta$ -neoendorphin | YGGFLRKYP |
| | α-neoendorphin | YGGFLRKYPK |
|  | Dyn A8 | YGGFLRRI |
|  | Dyn A13 | YGGFLRRIRPKLK |
|  | Dyn A17 | YGGFLRRIRPKLKWDNQ |
|  | Dyn B13 | YGGFLRRQFKVVT |
| Proenkephalin | Leu-enk | YGGFL |
|  | Met-enk | YGGFM |
|  | Met-enk RF | YGGFMRF |
|  | Met-enk RGL | YGGFMRGL |
|  | BAM12 | YGGFMRRVGRPE |
|  | BAM18 | YGGFMRRVGRPEWWMDYQ |
|  | BAM22 | YGGFMRRVGRPEWWMDYQKRYG |
|  | Peptide E | YGGFMRRVGRPEWWMDYQKRYGGFL |

### Co-localization of ECE2 and kappa opioid receptors

In order for ECE2 to be co-internalized to an endocytic compartment with KOR to regulate trafficking upon treatment with opioid peptide agonists, it would need to be expressed in close physical proximity to the receptor. To test this, we performed a proximity ligation assay (PLA) in recombinant CHO cells co-expressing Flag-tagged KOR and HA-tagged ECE2 treated with or without opioid peptide agonists (schematic in **Fig 2A**). We chose proDYN derived Dyn A17 and Dyn B13 as peptide agonists for our PLA studies since previous studies have shown treatment with these specific peptides sort KOR into different endosomal compartments (Kunselman et al., 2021). We find that in the absence of opioid peptide treatment, ECE2 and KOR are in close proximity at the cell surface and in intracellular compartments (**Fig 2B No drug).** Treatment with Dyn A17 and Dyn B13 (100 nM) for 10 minutes causes no significant changes in the extent of ECE2 and KOR co-localization (**Fig 2B Dyn A 10 min and Dyn B 10 min)**. However, 30 minutes of treatment with Dyn A17 but not Dyn B13 significantly increases the extent of ECE2 and KOR co-localization (**Fig 2B** Dyn A 30 min and Dyn B 30 min) (**Fig 2C ***p<0.001)**. Together, PLA studies show close proximity between KOR and ECE2 and that 30 min treatment with select proDYN derived peptides (i.e. Dyn A17) increases the extent of co-localization.

**Fig 2.**
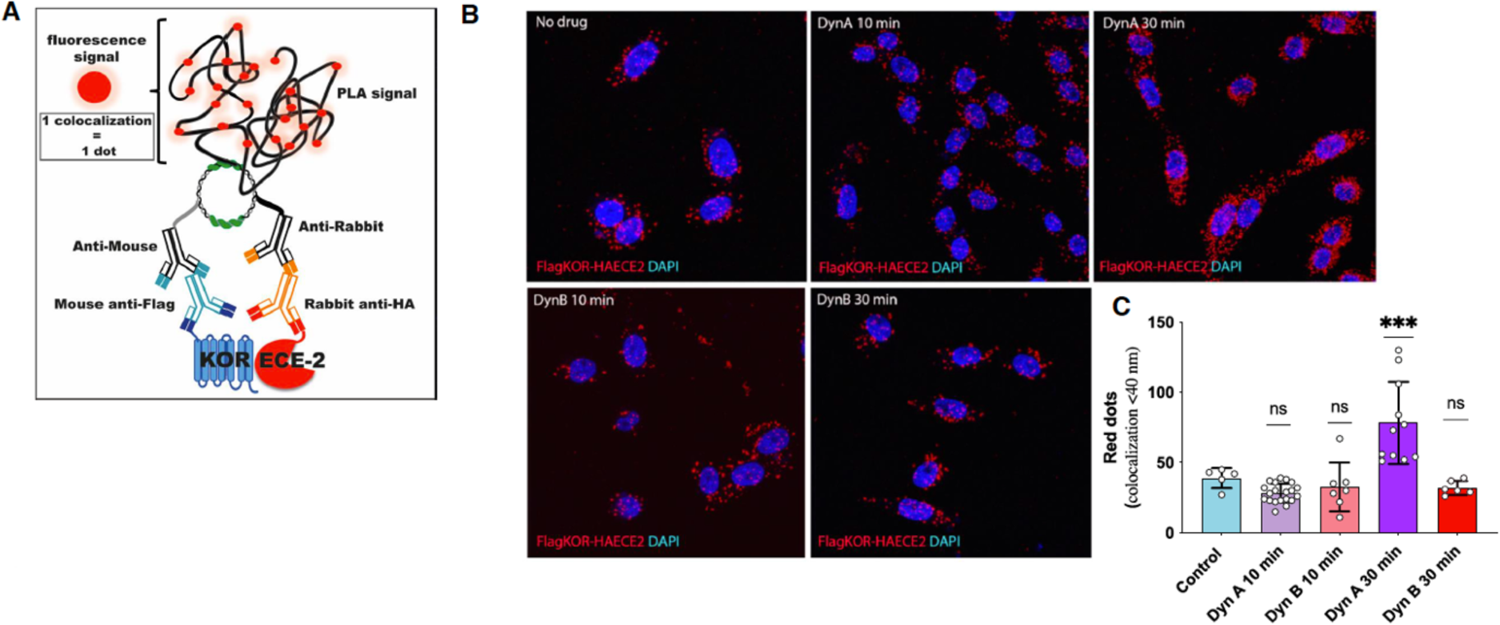
ECE2 co-localizes with KOR at the cell surface and intercellular compartments. **(A)** Schematic of proximity ligation assay to detect proximity between ECE2 and kappa opioid receptors (KOR). **(B)** Cells co-expressing Flag-tagged KOR and HA-tagged ECE2 were treated with vehicle (no drug; control), Dyn A17 or Dyn B13 for 10 or 30 min and co-localization of KOR with ECE2 examined using a proximity ligation assay as described in Methods. Individual red dots represent 1 colocalization event (less than 40 nm in proximity). Data represent Mean±SD. ***p<0.001; One-way ANOVA with Tukey’s multiple comparisons test.

### ECE2 internalization following KOR activation by proDYN-derived peptides

Next, we expanded our panel of proDYN-derived peptides to six physiologically relevant agonists (**Table 1**) and examined if KOR internalization caused by treatment with these peptides leads to internalization of ECE2 from the cell surface with a similar time course to that of KOR. Internalization studies were carried out using recombinant CHO cells co-expressing Flag-tagged KOR and HA-tagged ECE2. The extent of KOR and ECE2 internalization upon treatment with dynorphins was measured by assessing the time dependent loss of Flag-monoclonal antibody-labelled KOR and HA-polyclonal antibody-labelled ECE2 from the cell surface. We find that treatment with 100 nM of Dyn A8, Dyn A13, Dyn B13, Dyn A17, Alpha-neoendorphin or Beta-neoendorphin cause a decrease in cell surface expression of ECE2, with both the rate and extent of ECE2 internalization paralleling that of KOR (**Fig 3 A-F**). These results suggest that dynorphin peptides which show moderate differences in length and C-terminal peptide sequence induce similar levels of ECE2 and KOR internalization.

**Fig 3.**
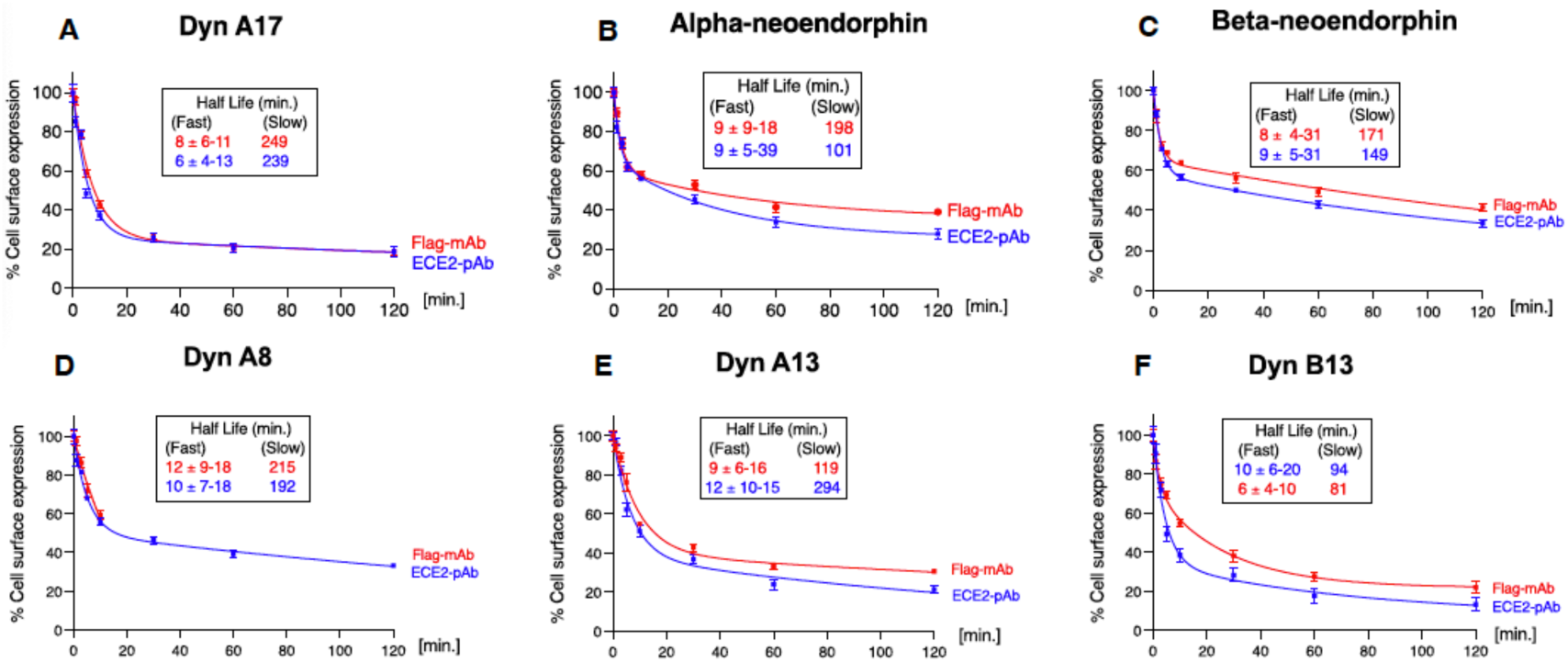
ECE2 internalization parallels the rate and extent of KOR internalization by proDYN derived peptides. CHO cells co-expressing FLAG-tagged KOR and HA-tagged ECE2 were prelabelled with primary antibodies to detect KOR and ECE2 at 4°C as described in Methods followed by treatment for 0-120 min at 37°C with 100 nM of either Dyn A17 **(A)**, Alpha-neoendorphin **(B)**, Beta-neoendorphin **(C)**, Dyn A8 (**D**), Dyn A13 **(E)**, or Dyn B13 **(F)** to induce receptor internalization and processed for ELISA as described in Methods. Cell surface levels of KOR and ECE2 at t=0 min was taken as 100%. Data represent Mean±SD n= 4 independent experiments in triplicate. Abbreviations: Flag mAB: Flag-monoclonal antibody-labelled KOR ECE2-pAb: HA-polyclonal antibody-labelled ECE2.

### ECE2 recycling following KOR internalization by proDYN-derived peptides

Next, we wondered if KOR and ECE2 show parallel recycling kinetics following treatment with the same proDYN-derived peptides; this is important since recycling plays a critical role in receptor resensitization (Hanyaloglu and Von Zastrow, 2008). To do so, CHO cells co-expressing FLAG-tagged KOR and HA-tagged ECE2 were treated with 100 nM of dynorphin peptides for 30 min to induce internalization, followed by incubation in media without the peptides to induce receptor recycling and measurement of the time dependent reappearance of cell surface Flag antibody-labelled KOR and antibody-labelled ECE2. We find that treatment with Dyn A17, Alpha-neoendorphin and Beta-neoendorphin causes ECE2 and KOR to recycle at a similar rate and extent (**Fig 4 A-C**). In contrast, treatment with Dyn A8, Dyn A13 and Dyn B13 (100 nM) causes recycling of KOR and ECE2 at differing rates and extent. For example, at 120 minutes 80% of KOR is recycled following treatment with Dyn B13 but only 58% of ECE2 is recycled (**Fig 4 D-F**). These results suggest that while treatment with proDYN-derived peptides do not affect the extent of internalization of KOR and ECE2, they differentially modulate recycling profiles.

**Fig 4.**
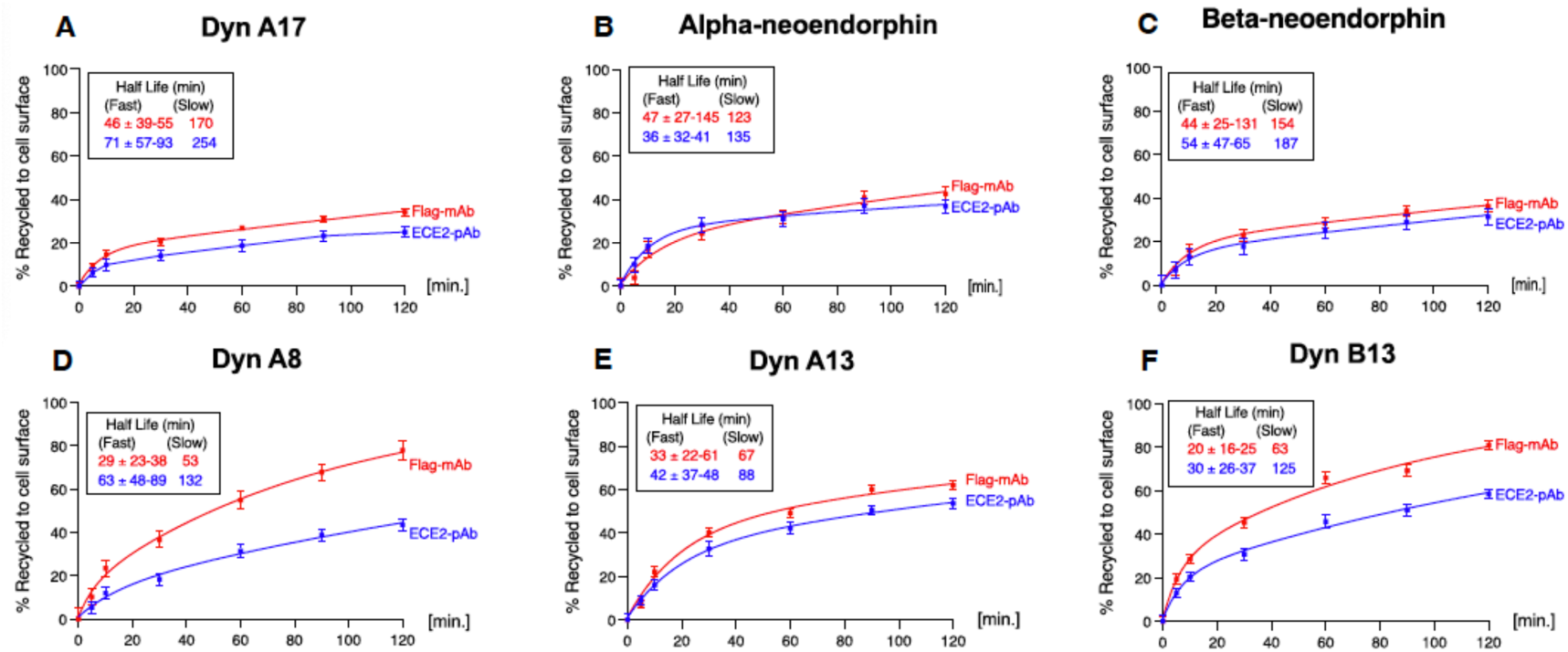
ECE2 recycling parallels the rate and extent of KOR recycling following internalization by proDYN derived peptides. CHO cells co-expressing FLAG-tagged KOR and HA-tagged ECE2 were treated with 100 nM of either Dyn A17 **(A)**, Alpha-neoendorphin **(B)**, Beta-neoendorphin **(C)**, Dyn A8 (**D**), Dyn A13 **(E)**, or Dyn B13 **(F)** for 30 minutes to induce internalization, followed by incubation in media without agonist for 0-120 min to induce receptor recycling, and then processed for ELISA as described in “Methods”. The percent recycled receptors are calculated by subtracting the percentage of surface receptors at t=0 (30 min internalization) from all time points. Data represent Mean±SD n= 4 independent experiments in triplicate.

### ECE2 internalization following KOR activation by proENK-derived peptides

Because proENK-derived peptides have recently been reported to bind KOR, but with varying affinities and abilities to induce bias signaling (Gomes et al., 2020), we next examined ECE2 and KOR internalization kinetics in response to treatment with eight physiologically relevant proENK-derived peptides (**Table 1**). We find that treatment with the shorter proENK-derived peptides Leu-enk, Met-enk, Met-enk RF or Met-enk RGL leads to lower levels of ECE2 and KOR internalization albeit with a comparable rate and extent of internalization (**Fig 5 A-D)**. In contrast, treatment with the longer peptides BAM 12, BAM 18, BAM 22, or Peptide E results in a more robust decrease in cell surface expression of ECE2 and KOR, and among them only treatment with Peptide E causes ECE2 to internalize at a rate and extent paralleling that of KOR (**Fig 5 E-H**). Taken together, these findings suggest that unlike proDYN derived peptides, the level of KOR and ECE2 internalization by proENK-derived peptides is differentially affected by different peptides.

**Fig 5.**
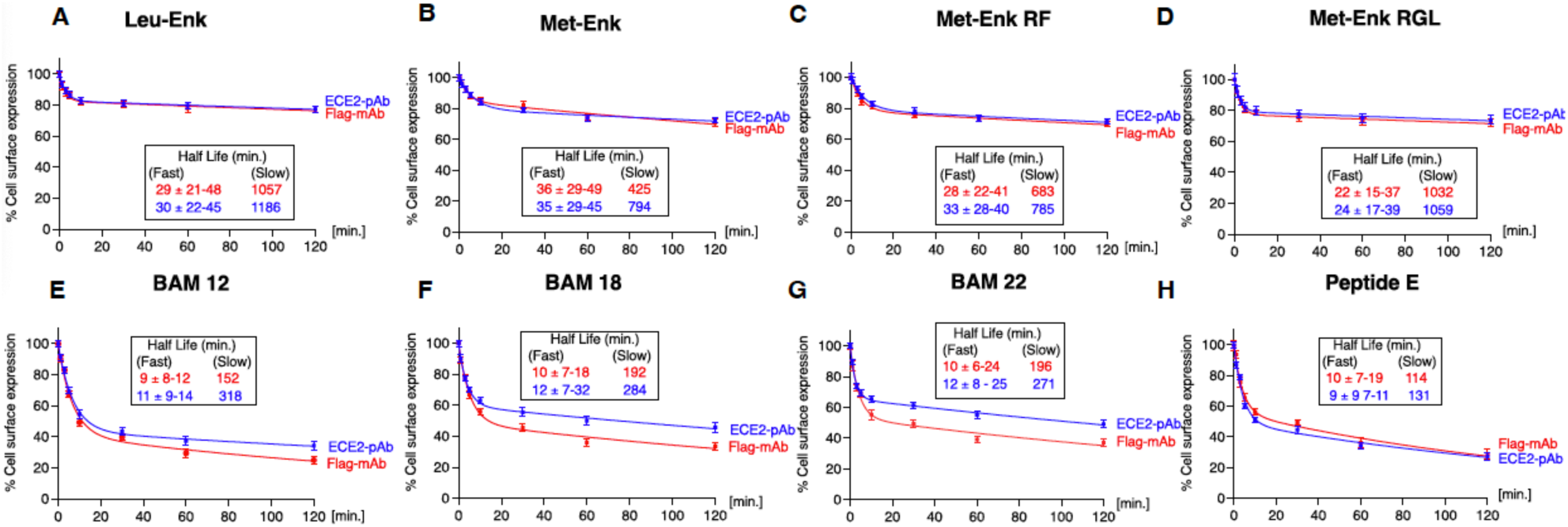
ECE2 internalization parallels the rate and extent of KOR internalization by proENK derived peptides. CHO cells co-expressing FLAG-tagged KOR and HA-tagged ECE2 were prelabelled with primary antibodies to detect KOR and ECE2 at 4°C as described in Methods followed by treatment for 0-120 min with 100 nM of either Leu-enkephalin **(A)**, Met-enkephalin **(B)**, Met-enkephalin RF **(C)**, Met-enkephalin RGL (**D**), BAM 12 **(E)**, BAM18 **(F)**, BAM22 **(G)** or Peptide E **(H)** to induce receptor internalization and processed for ELISA as described in Methods. Cell surface levels of KOR and ECE2 at t=0 min was taken as 100%. Mean±SD n= 4 independent experiments in triplicate.

### ECE2 recycling following KOR internalization by proENK-derived peptides

Next, we examined KOR and ECE2 recycling following internalization by proENK-derived peptides. We find that treatment with the shorter proENK-derived peptides, Leu-enk, Met-enk, Met-enk RF or Met-enk RGL leads to minimal recycling of ECE2, although the rate and extent generally parallels that of KOR (**Fig 6 A-D**). In contrast, treatment with the longer peptides BAM 12, BAM18, BAM 22 or Peptide E leads to more robust recycling of KOR and ECE2 (**Fig 6 E-H**). In addition, the extent of KOR and ECE2 recycling at 120 minutes differs between the longer proENK peptides, with BAM 22 causing more recycling of KOR (52%) compared to ECE2 (39%) and Peptide E causing less recycling of KOR (40%) compared to ECE2 (52%) (**Fig 6 E and H**). Together, these results indicate that levels of internalization induced by proENK-derived peptides determine the levels of recycling for KOR and ECE2 and that the rate and extent of recycling differ between different opioid peptides.

**Fig 6.**
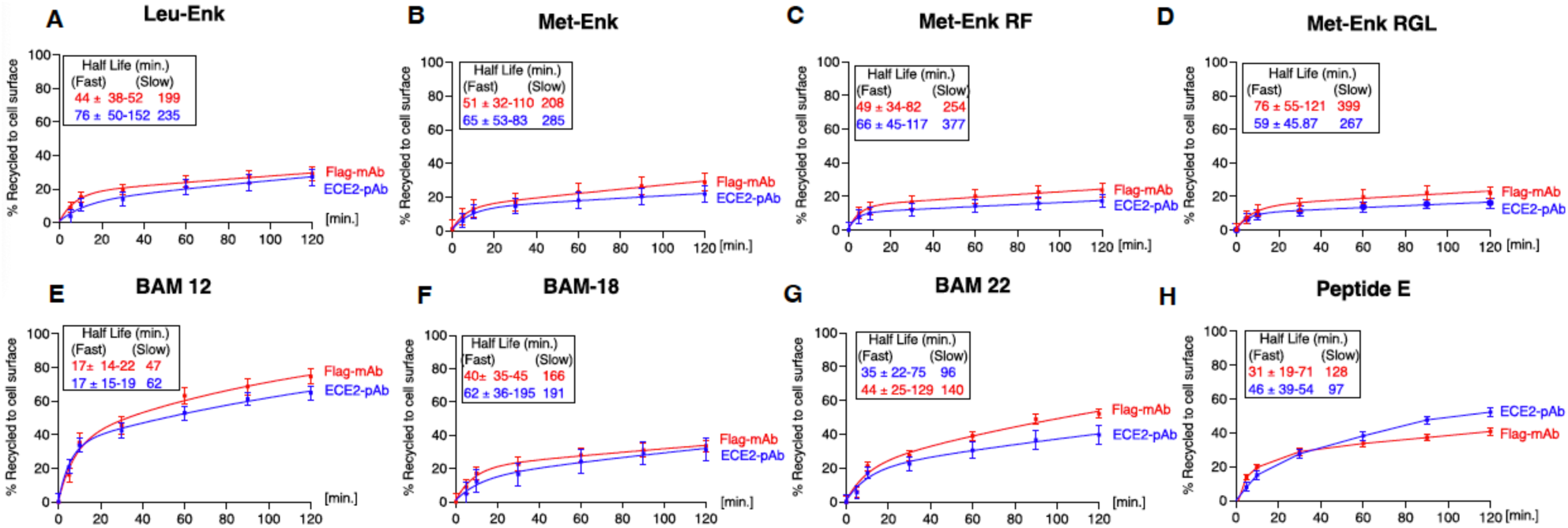
ECE2 recycling parallels the rate and extent of KOR recycling following internalization by proenkephalin derived peptides. CHO cells co-expressing FLAG-tagged KOR and HA-tagged ECE2 were treated with 100 nM of either Leu-enkephalin **(A)**, Met-enkephalin **(B)**, Met-enkephalin RF **(C)**, Met-enkephalin RGL (**D**), BAM 12 **(E)**, BAM18 **(F)**, BAM22 **(G)** or Peptide E **(H)** to induce receptor internalization, followed by incubation in media without agonist for 0-120 min to induce receptor recycling, and then processed for ELISA as described in “Methods”. The percent recycled receptors were calculated by subtracting the percentage of surface receptors at t=0 (30 min internalization) from all time points. Mean±SD n= 4 independent experiments in triplicate.

### KOR recycling is affected by the ECE2 inhibitor S136492

We previously showed that the rate and extent of MOR and DOR recycling but not internalization is regulated by ECE2 following treatment with opioid peptides that are its substrates (Gupta et al., 2014, 2015). Thus, we next asked if ECE2 substrate specificity could be playing a role in regulating KOR recycling. To do so, we used CHO cells co-expressing Flag-tagged KOR and HA-tagged ECE2 and examined the effect of the selective ECE2 inhibitor S136492 (Gagnidze et al., 2008) on KOR recycling. Cells were first treated with various KOR peptide agonists for 30 min to induce receptor internalization. The peptides were then removed to induce receptor recycling in the absence or presence of the ECE2 inhibitor for 60 min, and the reappearance of cell surface KOR was monitored by ELISA (**Schematic of protocol in Fig 7A**). We focused on recycling by the opioid peptide agonists Dyn B13 and Dyn A17 derived from proDYN and BAM-22, and Leu-enk derived from proENK due to differences in KOR recycling observed and because some of the peptides are ECE2 substrates and others are not (Mzhavia et al., 2003). We find that KOR recycling following internalization induced by Dyn B13, a known substrate of ECE2 (Mzhavia et al., 2003) is significantly reduced by the ECE2 inhibitor (**Fig 7B** left panel p<0.0001), while recycling by Dyn A17 (not a known substrate of ECE2 (Mzhavia et al., 2003) and a peptide agonist shown to traffic KOR to a degrative endocytic compartment (Kunselman et al., 2021) was not affected by the inhibitor (**Fig 7B** right panel). These results suggest that KOR recycling following internalization with proDYN-derived peptides is regulated by factors including the endocytic fate of the receptor and ECE2 substrate specificity.

**Fig 7.**
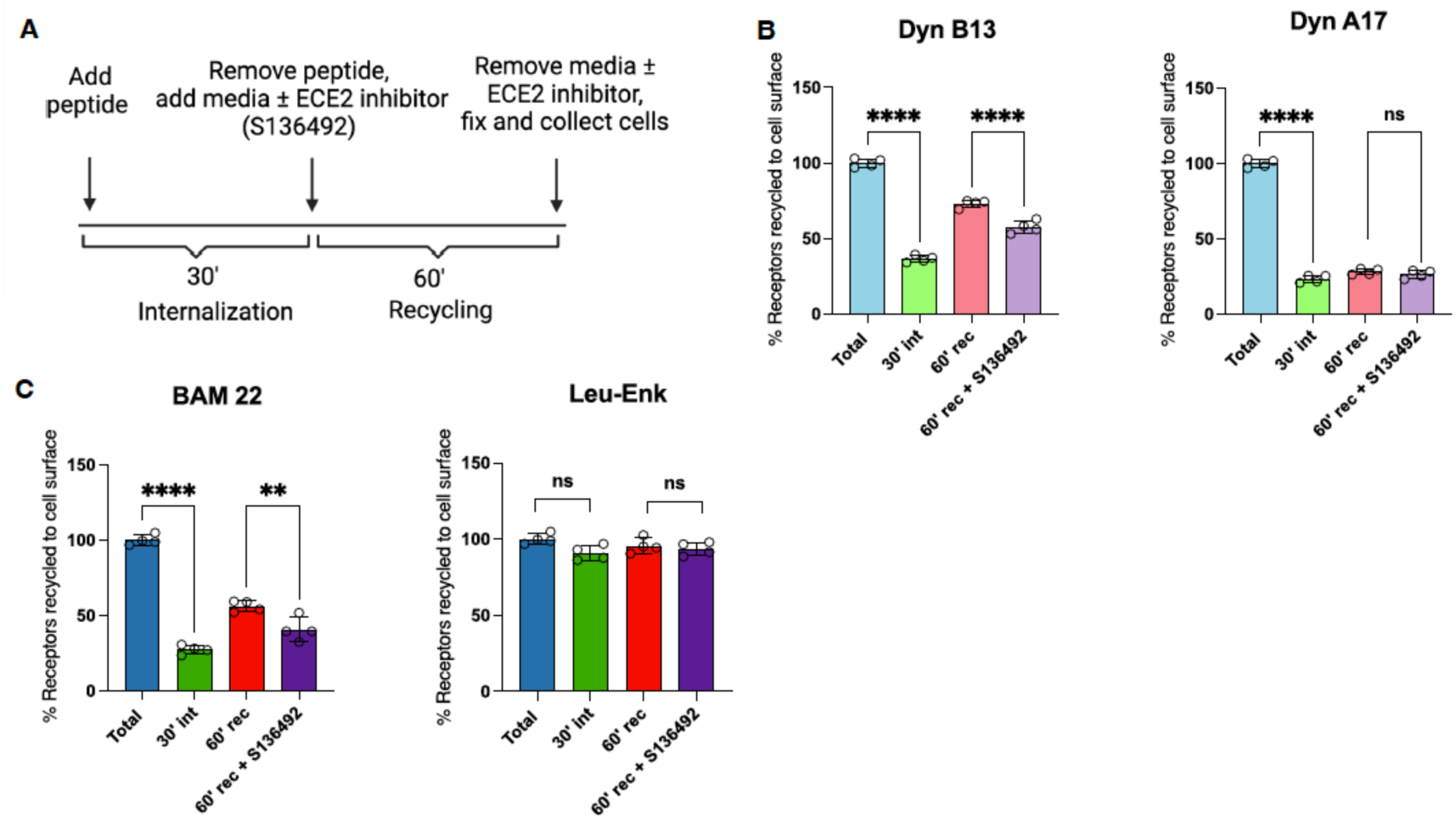
KOR recycling following internalization by opioid peptide substrates of ECE2 is reduced by the ECE2 inhibitor S136492. **(A)** Schematic of protocol to evaluate the effect of the ECE2 inhibitor, S136692 on KOR recycling. CHO cells co-expressing FLAG-tagged KOR and HA-tagged ECE2 were treated with 100 nM of peptides for 30 min to induce internalization (int), followed by incubation in media without peptides for 60 min to induce receptor recycling (rec) in the absence or presence of the ECE2 inhibitor, S136492, fixed and processed for ELISA as described in “Methods”. **(B)** Recycling after treatment with Dyn B13 but not Dyn A17 is reduced by ECE2 inhibition. **(C)** Recycling after treatment with BAM 22 but not Leu-enk is reduced by ECE-2 inhibition. The percent recycled receptors are calculated by subtracting the percentage of surface receptors at t=0 (30 min internalization) from all-time points. Mean±SD n= 4 independent experiments in triplicate. One-way ANOVA Tukey post-hoc *p < 0.05 **p < 0.01 ***p < 0.001 ****p < 0.0001.

Assessment of KOR recycling following internalization induced by proENK-derived peptides shows that the ECE2 inhibitor significantly reduces recycling by BAM 22 (**Fig 7C** left panel p<0.01) known to be a substrate of ECE2 (Mzhavia et al., 2003), whereas treatment with Leu-Enk, not a known ECE2 substrate, results in low levels of KOR recycling even in media without the ECE2 inhibitor (**Fig 7C** right panel). As treatment with Leu-enk, a low affinity KOR agonist (Gomes et al., 2020), also results in low KOR internalization (**Fig 5E**) these results suggest that KOR recycling following internalization by proENK derived peptides is influenced by the affinity of the peptide for the receptor and subsequent internalization patterns, and that ECE2 substrate specificity plays an additional role in proENK derived agonist mediated regulation of KOR recycling.

To assess the role of endogenous ECE2 activity in modulating KOR recycling, we used a DRG-derived cell line, F11, that expresses native ECE2 and opioid receptors (Fan et al., 1992; Gupta et al., 2015) (Schematic of protocol in **Fig 8A**). In line with our findings in CHO cells co-expressing KOR and ECE2, we find that KOR recycling following internalization by the ECE2 substrate, Dyn B13, is significantly reduced by the ECE2 inhibitor (**Fig 8B** left panel p=<0.001) while recycling following internalization induced by the non-substrate Dyn A17, is unaffected (**Fig 8B** right panel). In the case of the proENK-derived peptides, we find that recycling following internalization induced by BAM 22, a substrate of ECE2, is significantly reduced by the ECE2 inhibitor (**Fig 8C** left panel p=<0.0001) while recycling following internalization with Leu-enk, a low affinity KOR agonist and not a known substrate of ECE2, was unaffected (**Fig 8C** right panel). Together these results show that ECE2 plays a role in regulating KOR recycling by selectively processing peptide substrates following endocytosis in cells with endogenous expression of KOR and ECE2.

**Fig 8.**
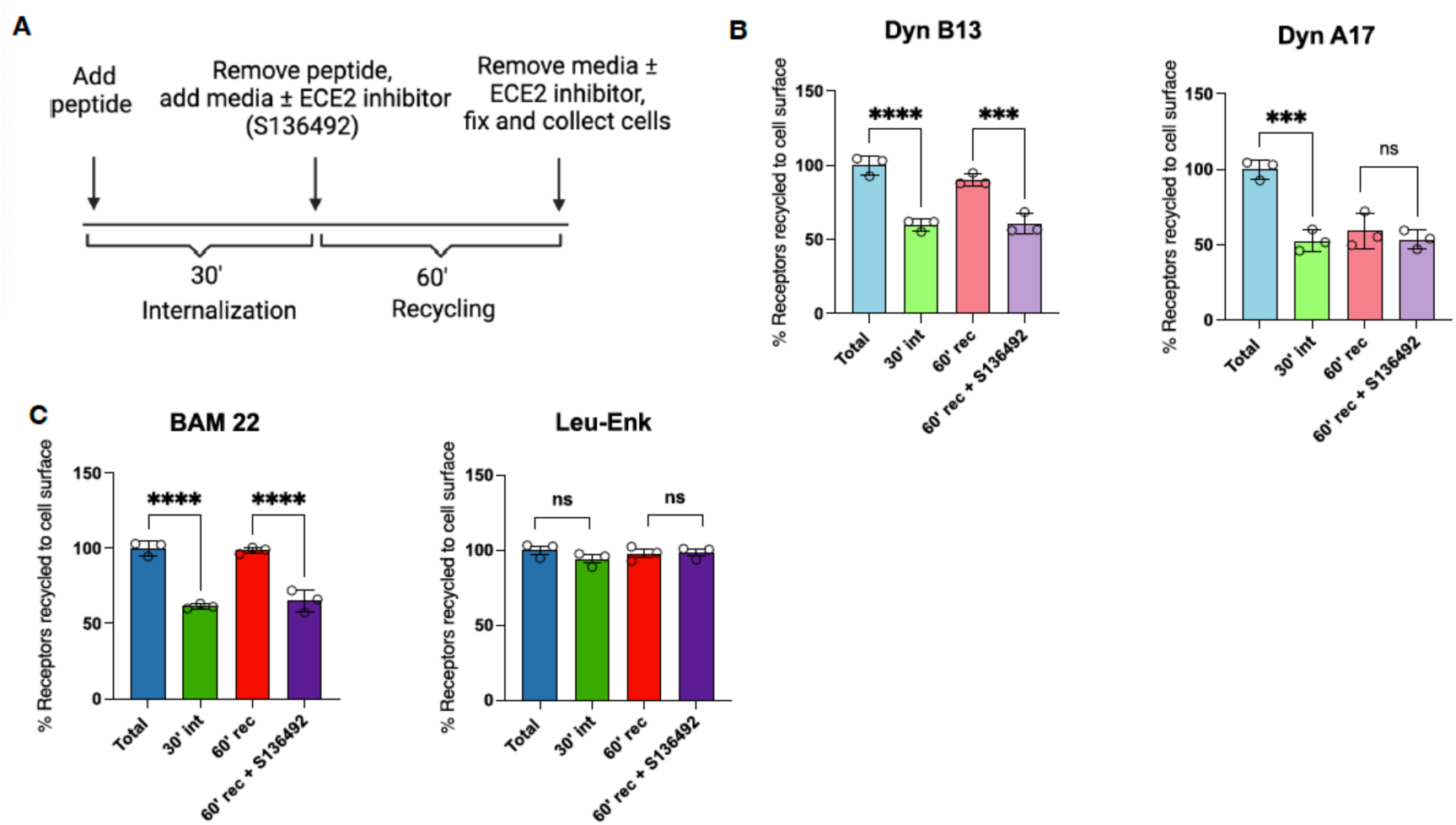
KOR recycling in F-11 cells following internalization by Dyn 13 and BAM 22 is reduced by the ECE2 inhibitor S136492. **(A)** Schematic of protocol to evaluate the effect of the ECE2 inhibitor, S136692 on KOR recycling in F-11 cells endogenously expressing KOR and ECE2. The cells were treated with 100 nM of peptides for 30 min to induce internalization (int), followed by incubation for 60 min in media without peptides and in the absence or presence of the ECE2 inhibitor, S136492, to induce receptor recycling (rec), fixed and KOR at the cell surface determined by ELISA as described in “Methods” **(B)** Recycling following internalization with Dyn B13 but not Dyn A17 is reduced by ECE2 inhibition. **(C)** Recycling following internalization with BAM 22 but not Leu-enk is reduced by ECE2 inhibition. The percent recycled receptors are calculated by subtracting the percentage of surface receptors at t=0 (30 min internalization) from all-time points. Mean±SD n= 4 independent experiments in triplicate. One-way ANOVA Tukey post-hoc *p < 0.05 **p < 0.01 ***p < 0.001 ****p < 0.0001.

### KOR signaling is affected by the ECE2 inhibitor S136492

Next, we examined the functional implications of ECE2 activity on KOR signaling by assessing the extent of signaling in the absence and presence of the ECE2 inhibitor, S136492. Since KOR activation leads to inhibition of adenylyl cyclase activity and subsequent reduction in cAMP levels, we used a cAMP assay to assess the effects of the ECE2 inhibitor on KOR signaling. For this, CHO cells co-expressing FLAG-tagged KOR and HA-tagged ECE2 were preincubated with forskolin to increase basal cAMP levels. This was followed by a 20 min treatment with peptides to induce receptor internalization. Peptides were removed and cells incubated in media in the absence or presence of 20 µM ECE2 inhibitor, S136492 for 60 min (mimicking conditions for KOR recycling). To assess receptor signal resensitization, the inhibitor was removed, and cells given a second pulse (5 min) with the peptide (Schematic of protocol in **Fig 9A**). We focused on signaling by the proDYN- and proENK-derived peptide known as substrates of ECE2, namely Dyn B13 and BAM 22. We find that in the absence of the ECE2 inhibitor, the second 5 min pulse of Dyn B13 and BAM 22 causes a decrease in cAMP levels indicating receptor signal resensitization following recycling (**Fig 9B-C light and dark purple bars respectively**). This decrease in cAMP levels is not observed in cells treated with the ECE2 inhibitor during the recycling phase (**Fig 9B-C light and dark pink bars respectively**). These results are consistent with ECE2 inhibition leading to retention of KOR in an intracellular compartment following treatment with opioid peptides that are ECE2 substrates, contributing to the loss of resensitization. Taken together these results suggest that by facilitating receptor recycling, ECE2 can modulate resensitization of KOR signaling by a specific subset of endogenous opioid peptides that serve as ECE2 substrates.

**Fig 9.**
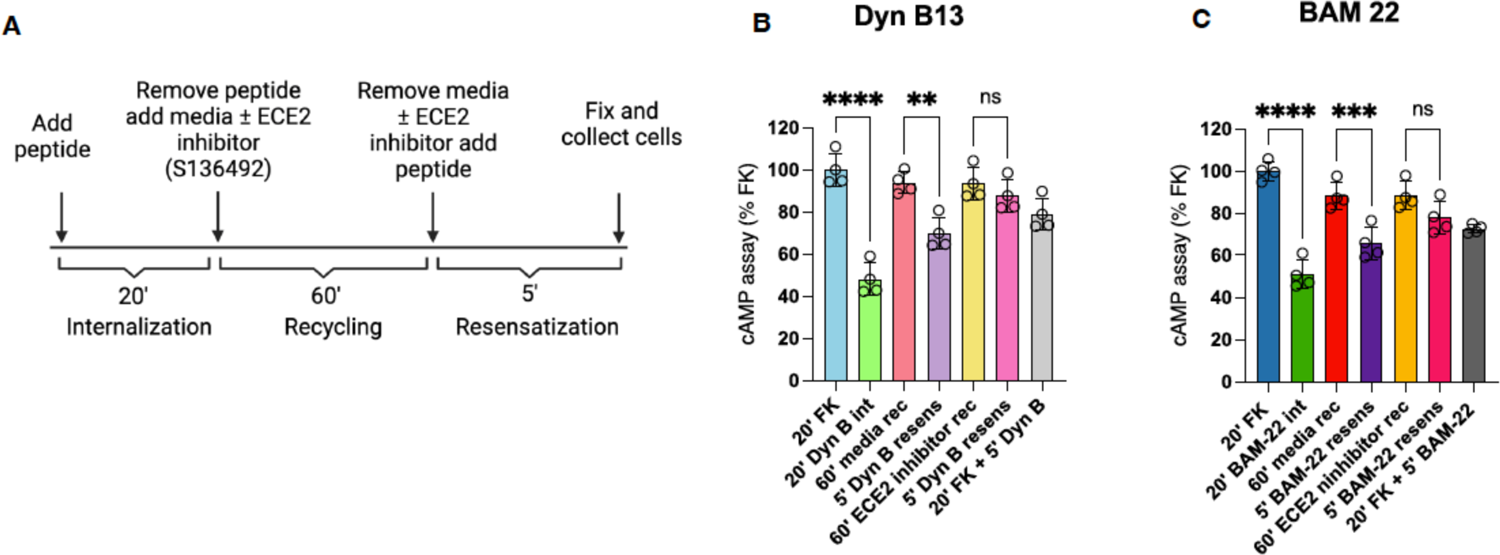
Signal resensitization by Dyn 13 and BAM 22 is blocked by the ECE2 inhibitor S136492. (**A**) CHO cells co-expressing FLAG-tagged KOR and HA-tagged ECE2 were treated with forskolin to increase basal cAMP levels, followed by a 20 min treatment with peptides. Peptides were removed, and cells were incubated in media in the absence or presence of 20 µM S136492 (ECE2 inhibitor) for 60 min. To assess receptor signal resensitization, the inhibitor was removed, and cells given a second pulse (5 min) of peptide. Signal resensitization by Dyn B13 **(B)** and BAM 22 **(C)** was blocked by ECE2 inhibitor. Mean±SD n= 4 independent experiments in triplicate. One-way ANOVA Tukey post-hoc *p < 0.05 **p < 0.01 ***p < 0.001 ****p < 0.0001.

## Discussion

This study investigated the role of ECE2 in regulating KOR function through co-internalization with the activated receptor and subsequent differential modulation of recycling and signaling. Using PLA, we demonstrate ECE2 expression at the cell surface in close proximity to KOR to facilitate co-internalization. Internalization and recycling studies further support co-trafficking, showing that ECE2 and KOR internalize and recycle at similar rates in an agonist-dependent manner. Using a selective ECE2 inhibitor, we demonstrate that ECE2 specifically modulates the recycling and resensitization of KOR in response to opioid peptide agonists that are its substrates. Taken with the differential expression of ECE2, proDYN and proENK mRNA in the brain, our findings suggest that ECE2 plays an important role in agonist mediated regulation of KOR function in select brain regions.

Studies examining the mRNA distribution of ECE2 in the brain show high expression in the dentate gyrus of the hippocampus, a brain region important for learning and memory, consistent with prior findings (Rodriguiz et al., 2008). ECE2 mRNA also overlaps with proDYN mRNA and proENK mRNA in this brain region, indicating its involvement in opioid peptide processing to mediate biological function. Indeed, mice lacking ECE2 have been shown to display deficits in learning and memory (Rodriguiz et al., 2008). Similarly, ECE2 has been shown to be highly expressed in the spinal cord (Miller et al., 2011), a key region in pain processing, and ECE2 KO mice show altered morphine responses (Miller et al., 2011). Together these findings suggest a role for ECE2 in processing opioid peptides involved in learning, memory, and pain. Importantly, moderate expression of ECE2 in the NAc but high levels of proDYN and proENK mRNA indicates that other metalloproteases could be playing a role in non-classical peptide processing in this brain region in addition to classical processing via prohormone convertases (Mzhavia et al., 2003).

While the cellular distribution of other metalloproteases including neprilysin and ECE1 are well characterized (Roosterman et al., 2007), ECE2 localization is less defined. Neprilysin is restricted to the cell surface, while ECE1 is found both at the cell surface and intracellular compartments (Roosterman et al., 2007). Our PLA studies reveal that ECE2 is expressed both at the cell surface and in intracellular compartments near KOR, and that Dyn A17 treatment increases KOR and ECE2 co-localization following 30 minutes of treatment. These findings are supported by our previous studies showing that ECE2 is in close proximity to DOR and MOR, both at the cell surface and in intracellular compartments and that treatment with peptide agonists Delt II and DAMGO, increases the extent of ECE2 co-localization with DOR and MOR, respectively (Gupta et al., 2014, 2015). Additionally, studies have shown that ECE2 localizes to both early and late endosomes (Pacheco-Quinto and Eckman, 2013; Gupta et al., 2015). Collectively these findings suggest that ECE2 is expressed at the cell surface near opioid receptors, where due to its acidic pH optimum it is inactive and following opioid receptor activation, it is endocytosed with the receptor-peptide complex into acidic endosomes to differentially regulate agonist-mediated receptor trafficking.

PLA studies further show that while 10 minute Dyn A17 and Dyn B13 treatment does not alter KOR-ECE2 co-localization, 30 minute Dyn A17 treatment significantly increases co-localization. This suggests that a proportion of KOR and ECE2 do not co-localize until 30 minutes of treatment and that ligand type affects trafficking. Indeed, KOR has been shown to locate to different endosomal compartments based on the dynorphin peptide that activates it (Kunselman et al., 2021). While Dyn A17 localizes KOR to the lysosomal pathway where it continues to signal intracellularly prior to degradation, Dyn B13 localizes KOR to recycling endosomes (Kunselman et al., 2021). This divergent post endocytic sorting of KOR could contribute to differences in co-localization seen 30 minutes following treatment with Dyn A17 and Dyn B13.

Further support for the agonist mediated regulation of KOR and ECE2 trafficking comes from our internalization and recycling studies. We show that while both Dyn B13 and Dyn A17 cause similar and robust extents of KOR and ECE2 internalization, Dyn A17 causes very little recycling of KOR and ECE2 at all time points assessed, consistent with Dyn A17 trafficking KOR to the lysosomal pathway for degradation, while Dyn B13 traffics KOR to recycling endosomes (Kunselman et al., 2021). Similarly, we found that while Dyn A8, Dyn A13, Alpha-neoendorphin and Beta-neoendorphin all caused similar and robust extents of KOR and ECE2 internalization, Alpha-neoendorphin and Beta-neoendorphin caused relatively lower levels of recycling, suggesting potential differences in post endocytic sorting of KOR-ECE2 complex by these peptides, which will be assessed in future studies. Differences in the post endocytic sorting and subsequent recycling of KOR and ECE2 following internalization induced by related proDYN-derived peptides often co-expressed in physiologically relevant brain regions (Nikoshkov et al., 2005; Corder et al., 2018), likely contributes to signaling diversity by otherwise seemingly redundant KOR agonists. Given that KOR has been shown to remain in an active conformation and continues to signal from late endosomes following activation by Dyn A17 (Kunselman et al., 2021), future studies assessing post endocytic sorting of KOR by Dyn A8, Dyn A13, Alpha-neoendorphin and Beta-neoendorphin should also assess spatial and temporal signaling of KOR by these peptides.

The mechanisms regulating internalization and divergent endosomal sorting of GPCRs are yet to be fully understood and have largely been studied using receptor mutants or depleting key components of the trafficking machinery (Hanyaloglu and Von Zastrow, 2008; Rosciglione et al., 2014). While a ligand’s affinity for a GPCR and ability to induce bias signaling via recruitment of ß-arrestin plays important roles in receptor internalization, these factors do not seem to fully account for downstream trafficking (Bagheri Tudashki et al., 2020). This is supported by our current data showing that proDYN peptides with similar high affinities for KOR and abilities to recruit ß-arrestin (Gomes et al., 2020), display drastic differences in subsequent recycling. Indeed, emerging evidence suggests that the ß-arrestin subtypes recruited to activated opioid receptors including ßarr2 (Audet et al., 2012; Charfi et al., 2018), play a role in ligand specific recycling dynamics. Moreover, as the post endocytic sorting of GPCRs involves interactions between the cytoplasmic tail of the receptor and trafficking machinery including endosomal-sorting complex required for transport (ESCRT), trafficking of KOR likely involves interactions with the PDZ interacting protein NHERF1/EBP50 (Liu-Chen, 2004). Other mechanisms implicated in agonist dependent trafficking of KOR include ubiquitination (Bowman and Puthenveedu, 2015) and other posttranslational modifications (Chiu et al., 2017; Mann et al., 2019).

In the case of proENK-derived peptides, both internalization and recycling of KOR and ECE2 were found to be agonist dependent. Treatment with the longer peptides BAM 12, BAM 18, BAM 22 and peptide E induced robust KOR and ECE2 internalization and recycling compared to the shorter peptides Leu-enk, Met-enk, Met-enk RF and Met-enk RGL. These results are in line with our recent studies showing that proENK-derived peptides are promiscuous and can bind to and activate KOR in addition to MOR and DOR, and further suggest that the extent of internalization and recycling of KOR is dependent on the amino acid sequence of the proENK derived peptide activating it. As the longer proENK-derived peptides have been shown to have higher affinities for KOR and recruitment of ß-arrestin, compared to the shorter peptides (Gomes et al., 2020), this suggests that a ligand’s affinity and ability to induce bias signaling may play important roles in internalization and subsequent recycling of KOR. Interestingly, the shorter peptides Leu-enk, Met-enk, Met-enk RF and Met-enk RGL have been shown to be less efficacious at recruiting β-arrestin to KOR compared to DOR and MOR (Gomes et al., 2020). Together internalization and recycling studies suggest that the peptide activating KOR likely locks the KOR-ECE2 complex into a specific conformation which allows or prevents interactions with the machinery involved in endocytosis and post-endocytic sorting, affecting internalization and recycling dynamics of the complex.

A key finding from this study is that recycling and subsequent KOR signaling by Dyn B13 and BAM22, known substrates of ECE2 (Mzhavia et al., 2003) was blocked by the ECE2 inhibitor S136492. This is in line with our previous findings showing that ECE2 inhibition prevents recycling of KOR by Dyn B13 (Kunselman et al., 2021) recycling of MOR by Dyn B13 (Gupta et al., 2015) and recycling of DOR by the synthetic peptide agonist Delt II, also a known substrate of ECE2 (Gupta et al., 2014). These results suggest that in addition to agonist mediated post endocytic sorting of KOR, ECE2 plays a crucial role by selectively processing opioid peptides and enabling receptor recycling and subsequent signaling.

In conclusion, our findings provide a novel approach to manipulating KOR cell surface levels and function via ECE2 inhibition, with clinical relevance for disorders where dysregulations in KOR signaling are implicated. Indeed, inhibitors of other metalloproteases have emerged as key players in messenger peptide regulation, with therapeutic implications in human disease (Fournié-Zaluski et al., 1994; Daull et al., 2005). Future studies should investigate the functional impact of ECE2 inhibition on KOR-related behaviors and disease models.

## List of abbreviations

BAM 12: bovine adrenal medulla 12
BAM 18: bovine adrenal medulla 18
BAM 22: bovine adrenal medulla 22
Dyn A17: dynorphin A17
Dyn A13: dynorphin 13
Dyn B13: dynorphin B13
Dyn A8: dynorphin A8
DOR: delta opioid receptor
ECE2: Endothelin converting enzyme 2
KOR: kappa opioid receptor
Leu-enk: leucine-enkephalin
MOR: mu opioid receptor
Met-enk: methionine-enkephalin
Met-enk RF: methionine-enkephalin Arg6Phe7
Met-enk RGL: methionine-enkephalin Arg6-Gly7-Leu8
NAc: nucleus accumbens
Peptide E: Peptide E
ProDYN: prodynorphin
ProENK: proenkephalin
PLA: proximity ligation assay.

## Acknowledgements

The authors declare that all the data supporting the findings of this study are contained within the paper. We thank Dr. D. Felsenfeld (Mount Sinai School of Medicine) for provision of F-11 cells.

## Authorship contributions

*Participated in research design:* IG, LAD *Conducted experiments:* AG

*Performed data analysis:* IG, AO

*Contributed to manuscript writing:* AO, IG, LAD

## Materials and Methods

### Materials

F12 media (cat. no. 11765-054), penicillin-streptomycin (cat. no. 15140-122), hygromycin (cat. no. 10687010), Lipofectamine 2000 (cat. no. 11668030), and polyclonal anti-ECE2 antibodies (cat. No. PA5-100495; RRID:AB_2850004) were from Thermo Fisher Scientific (Waltham, MA). FBS (cat. no. FBS-01) was from LDP, Inc. (Towaco, NJ). Geneticin (G418, cat. no. G-418-10) was from GoldBio (St. Louis, MO). Mouse anti-Flag M2 antibody (cat.no. F3165: RRID:AB_259529) and protease inhibitor cocktail (cat. no. P2714) were from Sigma-Aldrich (St. Louis, MO). Rabbit anti-HA antibody (cat. no. sc-805; RRID:AB_631618) was from Santa Cruz Biotechnology (Dallas, TX). Anti-mouse horseradish peroxidase (HRP, cat. no. PI-2000-1) and anti-rabbit horseradish peroxidase (HRP, cat. no. PI-1000-1; RRID:AB_2916034) were from Vector Laboratories (Newark, NJ). Alpha-neoendorphin (cat. no. 021-42), Beta-neoendorphin (cat. no. 021-44), Dyn A8 (cat. no. 021-10), Dyn A13 (cat. no. 021-21), Dyn A17 (cat. no. 021-03), Dyn B13 (cat. no. 021-37), BAM 12 (cat. no. 024-05), BAM 18 (cat. no. 024-06), BAM 22 (cat. no. 024-07), Leu-enk (cat. no. 024-21), Met-enk (cat. no. 024-35), Met-enk RF (cat. no. 024-50), Met-enk RGL (cat. no. 024-49), and peptide E (cat. no. 024-57) were from (Burlingame, CA). The PLA kit (cat. no. DUO92101) was from Millipore Sigma (Burlington, MA).The HitHunter cAMP HS chemiluminescence detection kit (cat. no. 90-0075LM2) was from DiscoveRx (Fremont, CA).

### Methods

#### Cell Culture and Transfection

CHO cells (ATCC cat. no. CRL-9618; RRID:CVCL_0214) expressing N-terminally FLAG epitope-tagged KOR were cultured in F12 medium supplemented with 10% FBS, penicillin-streptomycin, and 500 µg/ml Geneticin (G418). The cells were transfected with human HA epitope-tagged ECE2 using Lipofectamine 2000 following the manufacturer’s protocol (ThermoFisher), and clones with stable co-expression of KOR and ECE2 were selected with 500 µg/ml Geneticin and 250 µg/ml hygromycin B. F11 cells (Sigma Aldrich cat. no. 08062601) were cultured in F12 media containing 2 mM L-glutamine, 15% FBS, HAT supplement, and penicillin-streptomycin.

#### Proximity Ligation Assay (PLA)

CHO cells expressing Flag-tagged KOR and HA-tagged ECE2 were plated on a 16-well chamber slide (Thermo Scientific, Waltham, MA, USA). After 48 h, cells were treated with Dyn A17 or Dyn B13, (100 nM) for 30 mins. Cells were fixed in 4% paraformaldehyde for 15 min. Cells were washed with PBS containing 20 mM glycine to quench the aldehyde groups. Next, cells were permeabilized with PBS containing 0.1% Triton X-100 for 5 min. After permeabilization, PLA was performed using a commercially available kit (Millipore Sigma, Burlington, MA) as described elsewhere (Gomes et al., 2016). Primary antibody dilution was 1:500 for mouse anti-Flag M2 (Millipore-Sigma) and 1:500 for rabbit anti-HA (Santa Cruz, Dallas, TA, USA). Samples were observed under a Leica SP5 DMI laser-scanning confocal microscope using a 63 x /1.4 oil objective and sequential scanning with narrow band-pass filters (420-480 nm for DAPI and 560-615 nm for PLA signal). Analyze particle function from image J (NIH) was used to count particles as reported (Gomes et al., 2016).

#### Internalization Studies

CHO cells expressing Flag-tagged KOR and HA-tagged ECE2 (2 x 10^5^) were plated into each well of a 24-well poly-D-lysine-coated plate. The next day, cells were washed with 1X PBS (to remove FBS) and the levels of KOR and ECE2 at the cell surface determined by ELISA as described previously (Gupta et al., 2008, 2014). Briefly, cell surface KOR and ECE2 were prelabeled for 1 h at 4°C using a 1:500 dilution of anti-Flag M2 mouse monoclonal antibody for KOR or anti-ECE2 polyclonal antibody for ECE2. Cells were then treated for 0-120 min at 37^0^C with 100 nM of peptides in growth media containing protease inhibitor cocktail to induce KOR and ECE2 internalization. Cells were washed to remove the peptide agonist, fixed with chilled 4% paraformaldehyde (for 3 min), and processed for detection cell surface KOR and ECE2 using anti-mouse (1:500 dilution) and anti-rabbit (1:1000 dilution) antibodies coupled to horse-radish peroxidase as described previously (Gupta et al., 2008). The percent internalized receptors was calculated by taking cell surface levels of KOR and ECE2 at t=0 min as 100%.

#### Recycling Studies

Recycling studies were carried out with either CHO cells expressing Flag-tagged KOR and HA-tagged ECE2 or with F-11 cells essentially as described previously (Gupta et al., 2014). Briefly, KOR and ECE2 were internalized following pre-labelling of KOR with anti-Flag M2 mouse monoclonal antibody or ECE2 with anti-ECE2 polyclonal antibody by treatment with 100 nM of the peptides for 30 min as described above for Internalization studies. Cells were washed to remove the peptide agonist and incubated with media without the agonist for 0-120 min at 37°C to induce KOR and ECE2 recycling. At the end of the incubation, the medium was removed, cells were chilled to 4°C, and briefly fixed with 4% paraformaldehyde for 3 min; this fixation protocol allows the detection of only cell surface receptors but not intracellular receptors (Gupta et al., 2014). Cells were washed thrice with 1 x PBS (5 min each) and cell surface KOR and ECE2 levels detected as described above under Internalization studies. The percent recycled receptors was calculated by subtracting the percentage of surface receptors at t=0 (30 min internalization) from all time points. In a separate set of studies to examine the effect of the ECE2 inhibitor, S136492, on KOR recycling, the receptors were pre-labeled with anti-Flag M2 mouse monoclonal antibody as described above, treated for 30 min with peptide agonists to induce receptor internalization, and after removal of the peptide agonist, cells were incubated with media in the absence or presence of 20 μM ECE2 inhibitor S136492 for 60 mins at 37°C to block ECE2 activity during recycling.

#### Adenylyl cyclase activity assays

cAMP levels were determined as described previously (Gupta et al., 2014, 2015; Kunselman et al., 2021) with minor modifications. Briefly, CHO cells co-expressing FLAG-tagged Kor and HA-tagged ECE2 were seeded into a 96-well plate coated with poly-D-lysine. The cells were incubated with 100 nM Dyn B13 or BAM 22 for 20 minutes at 37°C in media containing 10 μM forskolin and 100 μM IBMX. Peptides were removed by washing one time with 100 µL of PBS, the cells were incubated in media with or without 20 µM S136492 (ECE2 inhibitor) for 60 minutes, and the level of signal resensitization assessed by exposing the cells to a second pulse (5 minutes) of peptide treatment. The product of adenylyl cyclase activity, cAMP, was determined under the various conditions using the HitHunter cAMP HS chemiluminescence detection kit from DiscoveRx.

#### Statistical Analysis

Data are expressed as Mean ± SD of 2-4 independent experiments in triplicate. Data were analyzed using One-Way ANOVA with Tukey’s or Dunnett’s multiple comparison test, with significance set at p< 0.05. All calculations were performed using GraphPad Prism 9 software (GraphPad Software, Inc., San Diego, CA, USA). Details on data analysis and normalization for individual data sets can be found in the Figure Legends. The ‘percent internalized’ was calculated by taking cell surface levels of KOR and ECE2 at t=0 min as 100%. To obtain the percent recycled KOR or ECE2, surface levels at t=0 (30 min agonist-mediated internalization) were taken to represent 0% recycled receptors and subtracted from each recycling time point.

